# Identification of CDRH3 Loops in the B Cell Receptor Repertoire that Can Be Engaged by Candidate Immunogens

**DOI:** 10.1101/2021.12.18.473225

**Authors:** Olivia Swanson, Joshua S. Martin Beem, Brianna Rhodes, Avivah Wang, Maggie Barr, Haiyan Chen, Robert Parks, Kevin O. Saunders, Barton F. Haynes, Kevin Wiehe, Mihai L. Azoitei

## Abstract

A major goal for the development of vaccines against rapidly mutating viruses, such as influenza or HIV, is to elicit antibodies with broad neutralization capacity. However, B cell precursors capable of maturing into broadly neutralizing antibodies (bnAbs) can be rare in the immune repertoire. Due to the stochastic nature of B cell receptor (BCR) rearrangement, a limited number of third heavy chain complementary determining region (CDRH3) sequences are identical between different individuals. Thus, in order to successfully engage broadly neutralizing antibody precursors that rely on their CDRH3 loop for antigen recognition, immunogens must be able to tolerate sequence diversity in the B cell receptor repertoire across an entire vaccinated population. Here, we present a combined experimental and computational approach to identify BCRs in the human repertoire with CDRH3 loops predicted to be engaged by a target immunogen. For a given antibody/antigen pair, deep mutational scanning was first used to measure the effect of CDRH3 loop substitution on binding. BCR sequences, isolated experimentally or generated *in silico*, were subsequently evaluated to identify CDRH3 loops expected to be bound by the candidate immunogen. We applied this method to characterize two HIV-1 germline-targeting immunogens and found differences in the frequencies with which they are expected to engage target B cells, thus illustrating how this approach can be used to evaluate candidate immunogens towards B cell precursors engagement and to inform immunogen optimization strategies for more effective vaccine design.

## Introduction

For an effective vaccine, it is essential that a candidate immunogen activates a large number of B cells from the natural immune repertoire (1). The efficiency of this process depends on the overall frequency of the target B cell population and the affinity of the immunogen for the respective B cell receptors (BCRs) (2, 3). Many broadly neutralizing antibodies (bnAbs), including ones that target HIV-1, influenza, and coronaviruses, have rare, long heavy chain CDR3 loops, which may limit the number of precursor B cells that can be engaged by vaccination for their elicitation **(Supplementary Table 1)**(4–14). For example, the bnAbs that target the HIV-1 V2 apex and V3 glycan epitopes on the viral Envelope typically contain CDRH3 loops of over 20 amino acids (15–21). Similarly, bnAbs that bind to the influenza neuraminidase or hemagglutinin utilize long CDRH3 loops for the majority of their viral contacts (7–9, 22). Because of their potency and broad recognition of diverse viral isolates, the elicitation of these types of antibodies is currently of interest as part of HIV-1 and universal influenza vaccine development efforts (11, 13, 23–30).

Current strategies to develop an HIV vaccine use isolated bnAbs as templates for immunogen design (11, 13, 14, 31–33). Critical to this approach is the development of “priming” immunogens that bind with high affinity to rare bnAb precursors that can subsequently mature into bnAbs. Multiple HIV bnAb lineages contain long CDRH3 loops of over 20 amino acids. Loops of this length typically have long segments generated by random N-nucleotide addition at the junction of germline V-D and D-J gene segments, which results in diverse CDRH3 loops across antibodies with the same immunogenetics (34, 35). Due to the stochastic nature of V(D)J recombination, there is limited overlap in the BCR repertoires of multiple subjects and identical CDRH3 sequences between different individuals are exceedingly rare (36, 37). Thus, for any priming immunogen to successful engage bnAb precursors across a vaccinee population, the immunogen must be able to bind to bnAb precursor BCRs with some CDRH3 sequence diversity.

Current strategies to identify natural BCRs that engage with a given immunogen use expensive and intricate experimental methods (33). In brief, the standard practice is to label human peripheral blood mononuclear cells (PBMCs) with the target molecule, and perform Fluorescence Activated Cell Sorting (FACS) for the desired phenotype followed by sequencing of BCRs from the isolated B cells. The resulting sequences are then produced recombinantly as IgG antibodies and characterized for binding to the target immunogen. This method is limited by the necessity to access and manipulate large numbers of human B cells. Even when such experiments are possible, this approach can be biased by the selection strategy and does not provide information to improve immunogen design for increased B cell precursor engagement.

To address these limitations, here we describe a mixed experimental and bioinformatic approach to identify BCRs in the natural human repertoire with CDRH3 loops predicted to be engaged by a target immunogen. For a given CDRH3 loop, our approach utilizes deep scanning mutagenesis to generate a substitution matrix that quantifies the effect of every possible single amino acid substitution to the binding of a candidate immunogen (**Figure 1**). The substitution matrix is then used to evaluate collections of BCRs isolated experimentally (34) or generated *in silico* (38), in order to identify CDRH3 loop sequences expected to be bound by the candidate immunogen. Here, we applied this platform to analyze the ability of two previously described HIV-1 vaccine candidates, CH505.M5.G458Y and 10.17DT, to engage precursors of broadly neutralizing antibodies against the conserved CD4 binding site and the glycan-V3 epitopes on the HIV-1 envelope (Env) protein. In animal models, both these immunogens have been shown to activate precursors of the HIV broadly neutralizing antibodies CH235.12 and DH270.6 respectively (11, 12). However, the potential of these SOSIP immunogens to engage human B cells from the natural repertoire, which is critical for their success as vaccines, remains uncharacterized. Using our platform (**Figure 1**), we found that the natural B cell repertoire contains a high number of BCRs with CDRH3 loop sequences that should permit engagement by CH505.M5.G458Y. In contrast, our analysis predicted that B cell activation by the 10.17DT immunogen will be more limited, and identified ways to optimize this immunogen for more robust precursor engagement. These results illustrate how our approach can inform vaccine development efforts towards eliciting broadly neutralizing antibodies against diverse viruses (11–13, 27, 28, 30, 39–41).

**Figure 1.**
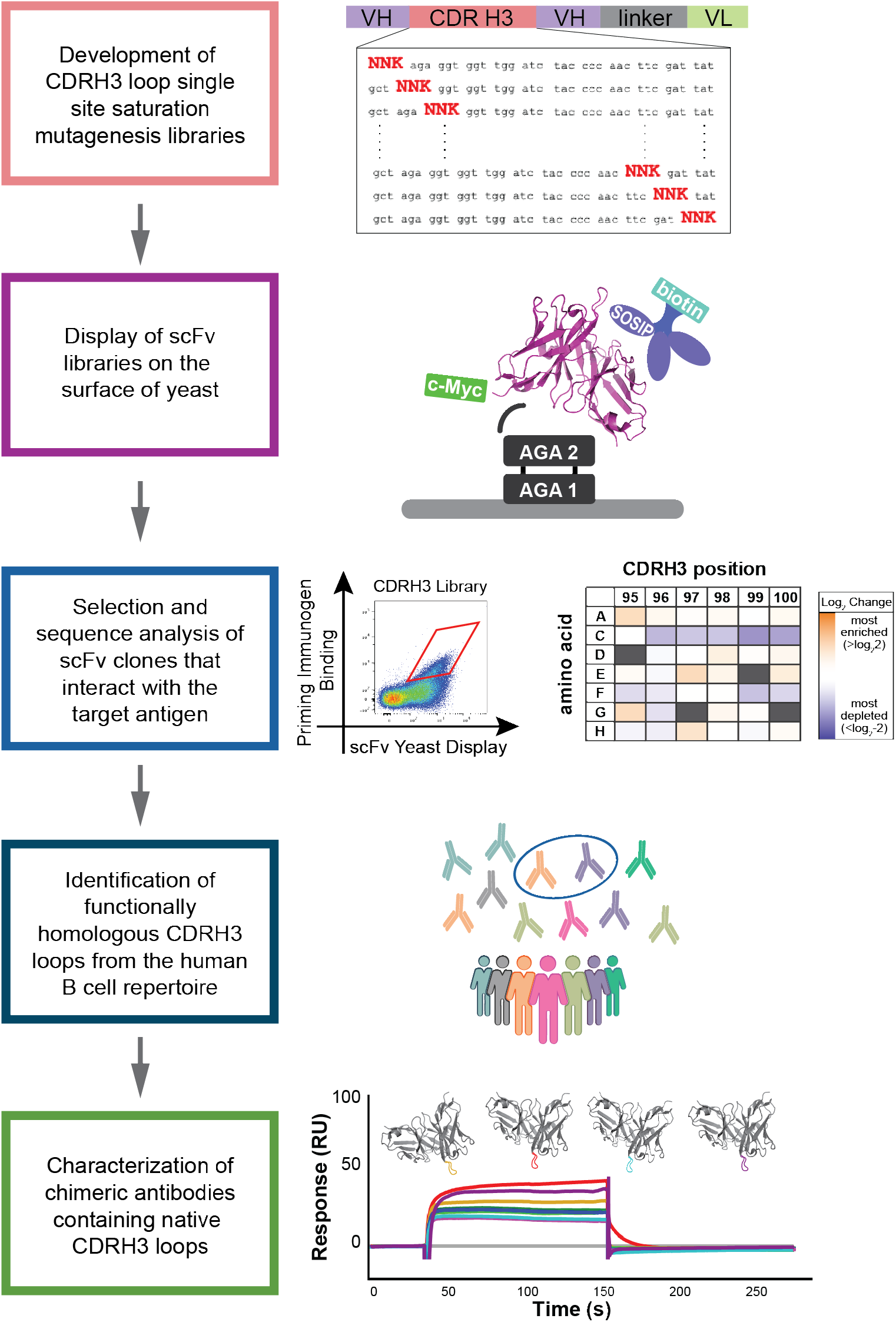
Combined experimental and bioinformatic workflow to characterize the ability of candidate immunogens towards the binding of diverse CDRH3 loops from the natural B Cell Receptor (BCR) repertoire.

## Results

### Identification of DH270UCA CDRH3 loop variants recognized by germline-targeting immunogen 10.17DT

The mature bnAb DH270.6 neutralizes an estimated 51% of circulating HIV-1 viruses by targeting the conserved glycan-V3 region on Env. The 10.17DT SOSIP was previously engineered to bind to the unmutated common ancestor (UCA) of DH270.6 with the goal of activating B cells precursors that can subsequently mature into bnAbs. Structural analysis revealed that the 10.17DT SOSIP makes strong interactions with the 20 amino acid CDRH3 loop of DH270 UCA. The CDRH3 loop contributes ~60% of the antibody buried surface in the 10.17DT binding complex (467Å^2^ out of 799Å^2^) and makes significant interactions with both the V3 loop as well as with the glycan present at position N332 (**Figure 2A, Supplementary Table 1**) (11). Given the extensive contacts between the 10.17DT SOSIP immunogen and the CDR H3 loop of DH270 UCA, it is likely that their interaction will be sensitive to the amino acid composition of the CDRH3 loop.

**Figure 2.**
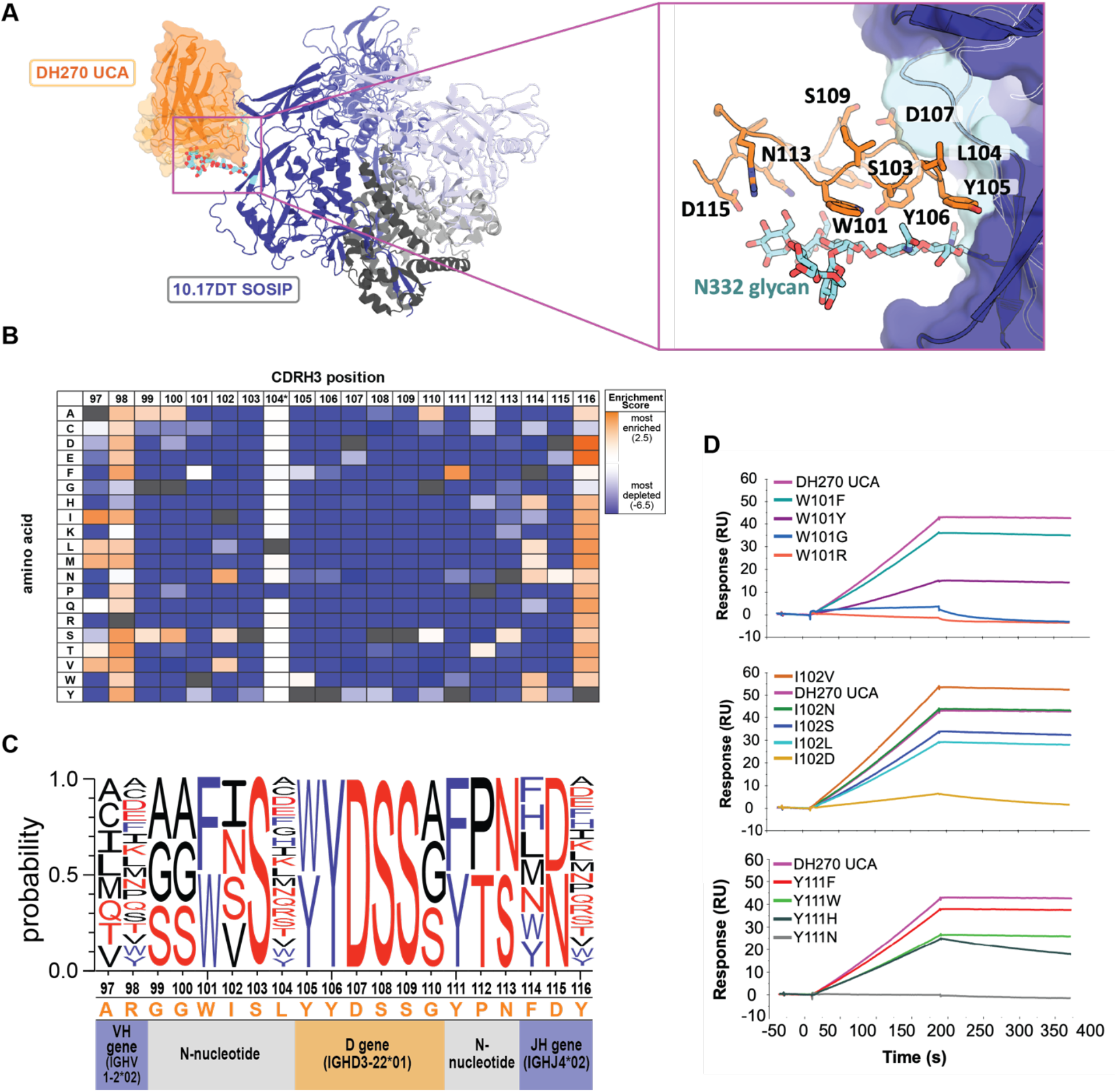
Immunogen recognition of CDRH3 loop substitutions in the unmutated common ancestor (UCA) of HIV-1 bnAb DH270. **A.** Structure of the 10.17DT germline targeting immunogen (*dark blue*) in complex with the DH270 UCA mAb (PDBid: 6um6; heavy chain: *orange*; light chain: *beige*). CDRH3 loop residues that interact with the glycan-V3 epitope (N332 glycan: *cyan sticks*; V3 loop base: *cyan surface*) are numbered and highlighted (*orange sticks*). **B.** Table of enrichment scores describing the effect of each amino acid substitution in the CDRH3 loop of DH270 UCA towards 10.17DT SOSIP binding. Values are colored from highly enriched (*red*) to significantly depleted (*blue*) and represent the average of two independent experiments.Native DH270 UCA CDRH3 residues are marked in black. Position 104 data was generated by ELISA. **C.** Logo plot depicting CDRH3 loop amino acid substitutions that maintained 10.17DT SOSIP binding (enrichment scores >−0.2). Non-polar residues are shown in *black*, polar residues in *red* and aromatic residues in *blue*. **D.** Binding of recombinant DH270 UCA IgGs containing single CDRH3 loop mutations to 10.17DT SOSIP.

To determine the ability of 10.17DT to engage antibody precursors with diverse CDRH3 loops, we first developed site saturation mutagenesis libraries that sampled all single amino acid substitutions in the CDRH3 loop of DH270 UCA. Libraries of scFv versions of DH270 UCA mutated antibodies were displayed on the surface of yeast and clones that maintained binding to 10.17DT were isolated by FACS (**Supplementary Figure 1**) (42). The DNA of selected cells was subsequently extracted and analyzed by next-generation sequencing. For a given CDRH3 substitution, its ability to engage 10.17DT was determined by an enrichment score calculated by taking the log_2_ of the frequency of the particular mutation in the 10.17DT sorted library divided by its frequency in the original, unsorted library. Enrichment scores above 0, corresponding to an increase in the frequency of a mutation upon sorting, indicated that the respective CDRH3 amino acid favors 10.17DT binding. Conversely, a negative enrichment score resulting from the depletion of CDRH3 variants in the 10.17DT selected clones, indicated that the corresponding amino acid substitution was detrimental to immunogen engagement. To ensure the reproducibility of this approach and to assess the consistency of the calculated enrichment value for a given mutation, the DH270 UCA CDRH3 library was evaluated for binding to 10.17DT in duplicate, by taking two aliquots of the unsorted library and analyzing them independently through all stages of the protocol. Individual enrichment scores were strongly correlated across the two replicates (**Supplementary Figure 2**). The correlation was strongest for enrichment scores higher than −5 (r^2^= 0.94) and decreased somewhat for values below this threshold (**Supplementary Figure 2**). An enrichment score of −5 corresponds to a 32-fold depletion of a particular mutation from the initial library upon sorting with 10.17DT. Highly depleted mutations are detected infrequently by FACS and their binding signal likely approaches that of background noise, which may explain the higher variation in their calculated enrichment scores across the two replicates. As will be shown below, enrichment scores below −3 correspond to a complete loss of 10.17DT binding for a given DH270 UCA CDRH3 mutation. Therefore, the actual magnitude of the enrichment levels for mutations that fall below the −3 level has limited biological significance. Taken together, these results demonstrate that the deep scanning mutagenesis platform robustly captures mutation enrichment levels in the range that captures diverse antigen binding outcomes.

Library screening and analysis revealed that a limited number of DH270 UCA CDRH3 loop single amino acids mutations maintained 10.17DT binding (**Figure 2B, 2C, Supplementary Figure 3**). Because limited diversity was observed in the naïve DH270 UCA CDRH3 loop library at position 104 upon sequencing, the effect of substitution at this site on 10.17DT binding was further assessed using a standard approach; IgG antibody variants containing all 19 single amino acid substitutions were expressed recombinantly and their ELISA binding to 10.17DT was compared to that of the native DH270 UCA (**Supplementary Figure 4**). The CDRH3 loop of DH270 UCA is encoded by VH1-2*02, D3-22*01, and JH4*02 genes, with 9 amino acids encoded by non-templated N-nucleotide additions. On average, only 5.3 different amino acids were tolerated by 10.17DT at each position in the CDRH3. The largest number of functional variants was found in the V_H_ and J_H_ templated regions, where only one site tolerated fewer than six different amino acids. The composition of the D-gene and N-addition encoded residues was more restricted, with an average of 3.5 different amino acid substitutions that maintained immunogen binding identified at each site. These data imply that 10.17DT will only engage natural BCRs containing CDRH3 loops that are highly similar in sequence to that of DH270 UCA.

To experimentally validate the results of scFv library screening, 16 DH270 UCA antibodies mutated at positions 101, 103, 104, 111 or 113 were expressed as recombinant IgG proteins and tested for binding to purified 10.17DT SOSIP by surface plasmon resonance (SPR) (**Figure 2D**). The binding levels of the four mAbs that contained mutations at position 101 closely mirrored the enrichment scores determined by deep scanning mutagenesis. Mutations W101R and W100G (enrichment scores of −2.6) caused almost complete loss of 10.17DT immunogen binding. W101F (enrichment score= 0) showed similar binding to the WT antibody while W101Y (enrichment score= −0.8) showed reduced binding, but not to the level of W101R or W101G. At position 102, mutations to Leu (enrichment score= −1.1) and Asp (enrichment score= −4.8) reduced binding, while changes to Asn (enrichment score= 1.7) and Val (enrichment score= 1) maintained or improved the 10.17DT interactions. A discrepancy was observed for the L102S mutation which was predicted to be favorable for binding by deep scanning mutagenesis but caused a slight reduction in the affinity of the purified IgG. At position 111, three mutations predicted to reduce binding by screening (Y111W, Y111H, Y111N) did so as recombinant IgGs, while another change (Y111F) maintained tight interactions as expected, capturing the trend, but not necessarily the magnitude, of their computed enrichment values. Finally, mutations were introduced at positions 103 and 113 that replaced the WT amino acids with closely related ones that were disfavored by deep scanning mutagenesis, to confirm the potential loss of binding from what appeared to be conservative mutations. Indeed, experimentally characterized DH270 UCA IgGs containing the S103A, S103T, or N113D mutations showed significantly reduced binding to 10.17DT (**Supplementary Figure 5**). These data illustrate that the deep scanning mutagenesis analysis of the DH270 UCA CDRH3 loop adequately recapitulates the affinity trends of recombinant IgGs containing the corresponding single amino acid changes. Binding discrepancies between deep mutational libraries and recombinant IgGs may occasionally occur as we have observed here and others reported previously (43, 44), due to the two different platforms used to measure binding that employ distinct antibody formats (FACS of scFvs displayed on the surface of yeast versus SPR of recombinant IgGs).

### Identification of CDRH3 loop variants recognized by germline-targeting immunogens that bind CH235 bnAb precursors

A similar deep scanning mutagenesis analysis was performed to identify the CH235 UCA CDRH3 loop variants that can be engaged by the germline targeting SOSIP immunogen CH505.M5.G458Y. Unlike DH270 UCA, the CDRH3 loop of CH235 UCA does not play a major role in antigen recognition. In the structure of the CH235 UCA-CH505.M5.G458Y complex, the 13 amino acid CDRH3 loop contributes only ~30% (245Å^2^ out of 858Å^2^) of the total antibody buried surface at the interface (**Figure 3A, Supplementary Table 1**) (12). It was therefore expected that CH505.M5.G458Y would recognize a high number of CH235 UCA CDRH3 loop variants.

**Figure 3.**
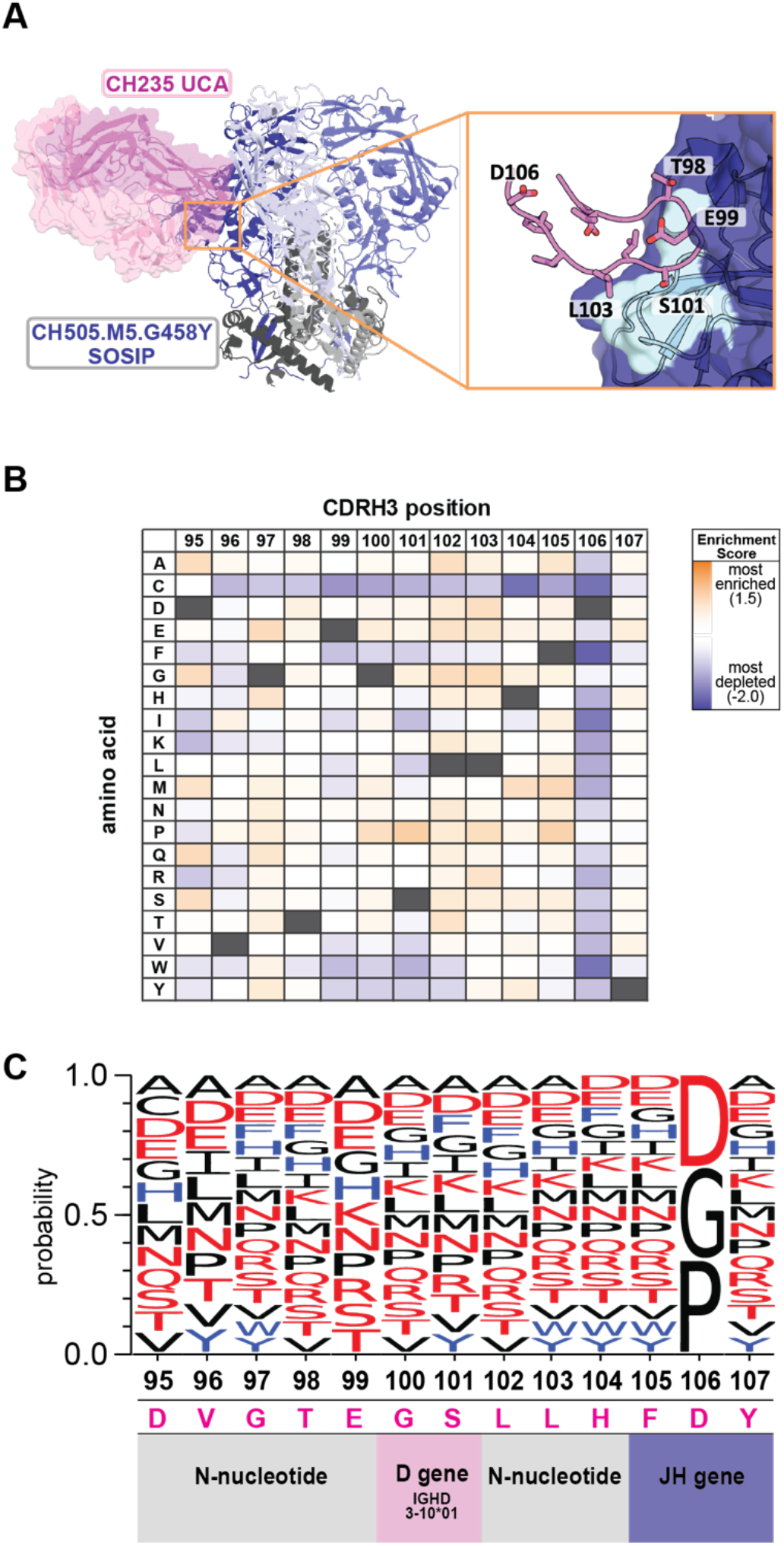
Immunogen recognition of CDRH3 loop substitutions in the unmutated common ancestor (UCA) of HIV-1 bnAb CH235. **A.** Structure of the CH505.M5.G458Y SOSIP germline targeting immunogen (*dark blue*) in complex with CH235 UCA mAb (PDBid: 6uda; heavy chain: *magenta*; light chain: *light pink*). CDRH3 loop residues that directly contact the CD4 binding site epitope (*cyan*) are highlighted and numbered (*magenta sticks*) **B.** Table of log_2_ enrichment ratios describing the effect of each amino acid substitution in the CDRH3 loop of CH235 UCA towards CH505.M5.G458Y SOSIP binding. Values are colored from highly enriched (*red*) to significantly depleted (*blue*) and represent the average of two independent experiments.Native CDRH3 residues are marked in black. **C.** Logo plot depicting CH235 UCA CDRH3 loop amino acid substitutions that maintained CH505.M5.G458Y SOSIP binding (enrichment score >−0.2). Non-polar residues are shown in *black*, polar residues in *red* and aromatic residues in *blue*.

Indeed, CH505.M5.G458Y maintained significant binding to a large number of CDRH3 substitutions in the CH235 UCA (**Figure 3B, C, Supplementary Figure 3A**). The 13 residue CDRH3 loop of CH235 UCA is encoded by the VH1-46, D3-10*01 and JH4*02 genes, with eight residues inserted by N-nucleotide additions. At the 10 positions encompassed by the N-nucleotide and D gene regions, the immunogen recognized an average of 15.3 amino acids. These results indicate that CH505.M5.G458Y engagement of BCRs upon vaccination will not be restricted by the amino acid composition of their CDRH3 loops, since this immunogen can bind CH235 UCA variants with highly divergent sequences in this region.

### Identification of CDRH3 loops in the human BCR repertoire that can be bound by 10.17DT and CH505.M5.G458Y

We next sought to identify CDRH3 loops in the human BCR repertoire that can be engaged by 10.17DT or CH505.M5.G458Y SOSIPs as a way to estimate the number of B cells that these molecules could engage upon vaccination. To find similar CDRH3 sequences to those recognized by these immunogens, we searched a human BCR database (36) that contains ~85 million non-redundant functional BCR sequences isolated from 10 individuals. In order to determine the frequency of DH270-like CDRH3s, we selected CDRH3 loops that had the same length (20 amino acids) as DH270 UCA and that contained the “YDSS” sequence at position 10 corresponding to the D gene (D3-22) of DH270 UCA. Of the 85,149,053 sequences analyzed, 47,975 CDRH3 loops matched these criteria, yielding an estimated frequency of 1 in 1,774 of BCRs with DH270-like CDRH3 loops. Next, we aligned these DH270-like CDRH3s against the CDRH3 substitution profile (**Figure 2C**) from the deep scanning mutagenesis to predict which CDRH3s contain only amino acids compatible with 10.17DT binding. Based on the experimental data (**Figure 2B, D**), any amino acid in the CDRH3 substitution profile with a log enrichment score below −0.2 upon 10.17DT selection was considered highly detrimental to 10.17DT binding and counted as a mismatch in the alignments. Of the 47,975 DH270-like CDRH3s analyzed, no sequences matched the amino acid variants in the substitution profile at all CDRH3 positions, and only one sequence contained just one mismatch (**Supplementary Figure 6A**). Therefore, the predicted frequency of BCRs expected to be bound by 10.17DT with similar high affinity as that for DH270 UCA (*K*_D_=507nM) is below the 1 in 85 million limit of detection set by the size of the analyzed database (11).

Next, we ranked the CDRH3 loops from the database for their ability to interact with 10.17DT based on the deep scanning mutagenesis data. This was done to assess experimentally the ability of 10.17DT to bind human DH270 precursors that are closely related, but not identical, to DH270 UCA and further validate the results of the deep scanning mutagenesis analysis. All the CDRH3 loops in the database with immunogenetics that matched those of DH270 UCA were scored based on their sequence similarity to the DH270 UCA CDRH3 substitution profile (**Supplementary Table 2**). The top 100 ranked antibodies were expressed as scFvs and displayed together on the surface of yeast. Upon sorting for two rounds with 10.17DT, no high-affinity binding was observed by FACS, although some scFv sequences were enriched in the selected clones, indicative of low affinity for the antigen (data not shown). Six such chimeric antibodies were expressed and purified as recombinant IgG and their binding to 10.17DT was measured by SPR. 10.17DT SOSIP bound weakly to these DH270 UCA chimeric antibodies containing natural CDRH3 loops, with binding levels less than 10% of that measured for the unmutated DH270 UCA mAb (**Figure 4, Supplementary Figure 7**), which precluded the determination of dissociation constants. These weak affinities were expected since all the CDRH3 loops tested experimentally contained at least two amino acids that were found by deep scanning mutagenesis to significantly reduce 10.17DT binding.

**Figure 4.**
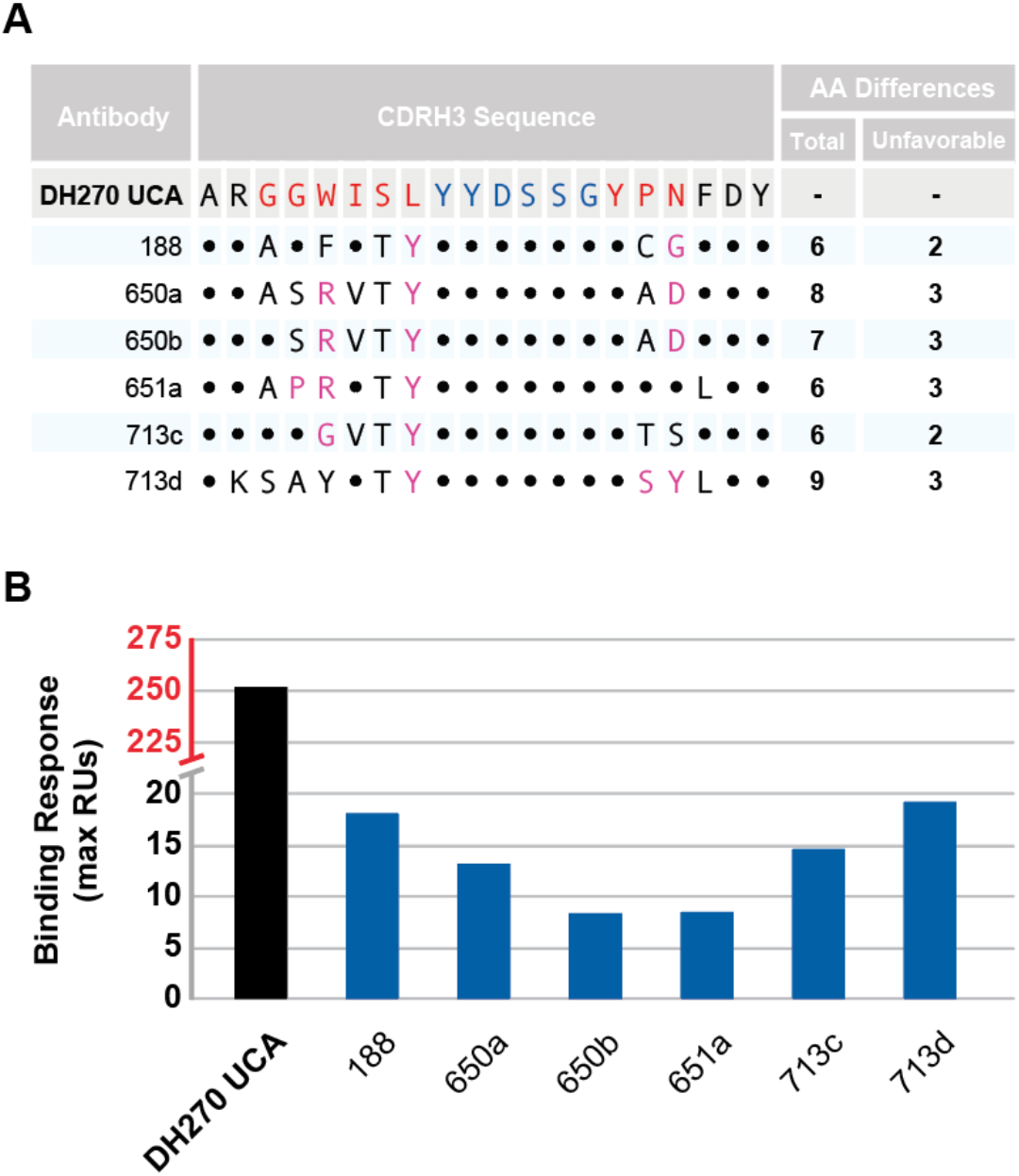
10.17DT SOSIP recognition of natural CDRH3 loops. **A.** Sequence alignment between the CDRH3 loop of DH270UCA and seven natural CDRH3 loops selected from a database of human BCRs and predicted to be recognized by 10.17DT SOSIP. Sequence changes expected to be neutral (*black*) and detrimental (*magenta*) towards 10.17DT SOSIP binding are indicated. DH270 UCA CDRH3 residues encoded by the D gene are shown in *blue*, while those encoded by N-nucleotides are in *red*. **B.** Binding of DH270UCA chimeric mAbs with the CDRH3 loop replaced by those described in (**A**) to 10.17DT SOSIP.

Because the database of human BCRs contains only a small fraction of the diversity expected to be present in the immune repertoire of an individual at any one time (estimated to be 10^9^-10^12^ unique BCRs) (36, 45, 46), we next wanted to determine if CDRH3s predicted to be compatible with 10.17DT binding could be present in a set of sequences that was the same order of magnitude in size as the human B cell repertoire. To address the limitation of available repertoire depth in the BCR database, we used the computational program IGoR (38) to simulate the V(D)J recombination process and to randomly generate ~2.5 trillion CDRH3 sequences. Amino acid frequencies at each position in the simulated CDRH3s were remarkably similar to those observed in the natural BCR sequence database despite the large difference in the total number of sequences (~48,000 versus 1.15 billion) (**Figure 5A)**. Next, we generated heatmaps for both sets of sequences by binning the CDRH3 loops according to the number of amino acids in each CDRH3 sequence that are incompatible with 10.17DT binding based on the deep scanning mutagenesis profile (**Figure 2B**) at various enrichment score threshold values (**Figure 5B**). A comparison of the heatmaps for the simulated and natural CDRH3 sequences revealed remarkably similar distributions of natural CDRH3 sequences and those simulated by IGoR. IGoR produced a smoother histogram due to the detection of sequences expected to bind 10.17DT with high affinity in the 1 in 10 million to 1 in 1 billion frequency range, a sampling level that exceeded the sequencing depth in the BCR database. Taken together, these data indicate that sampling from IGoR-simulated sequences can recapitulate the distribution of CDRH3s in the human naïve BCR repertoire and allows for the observation of extremely rare V(D)J recombination events.

**Figure 5.**
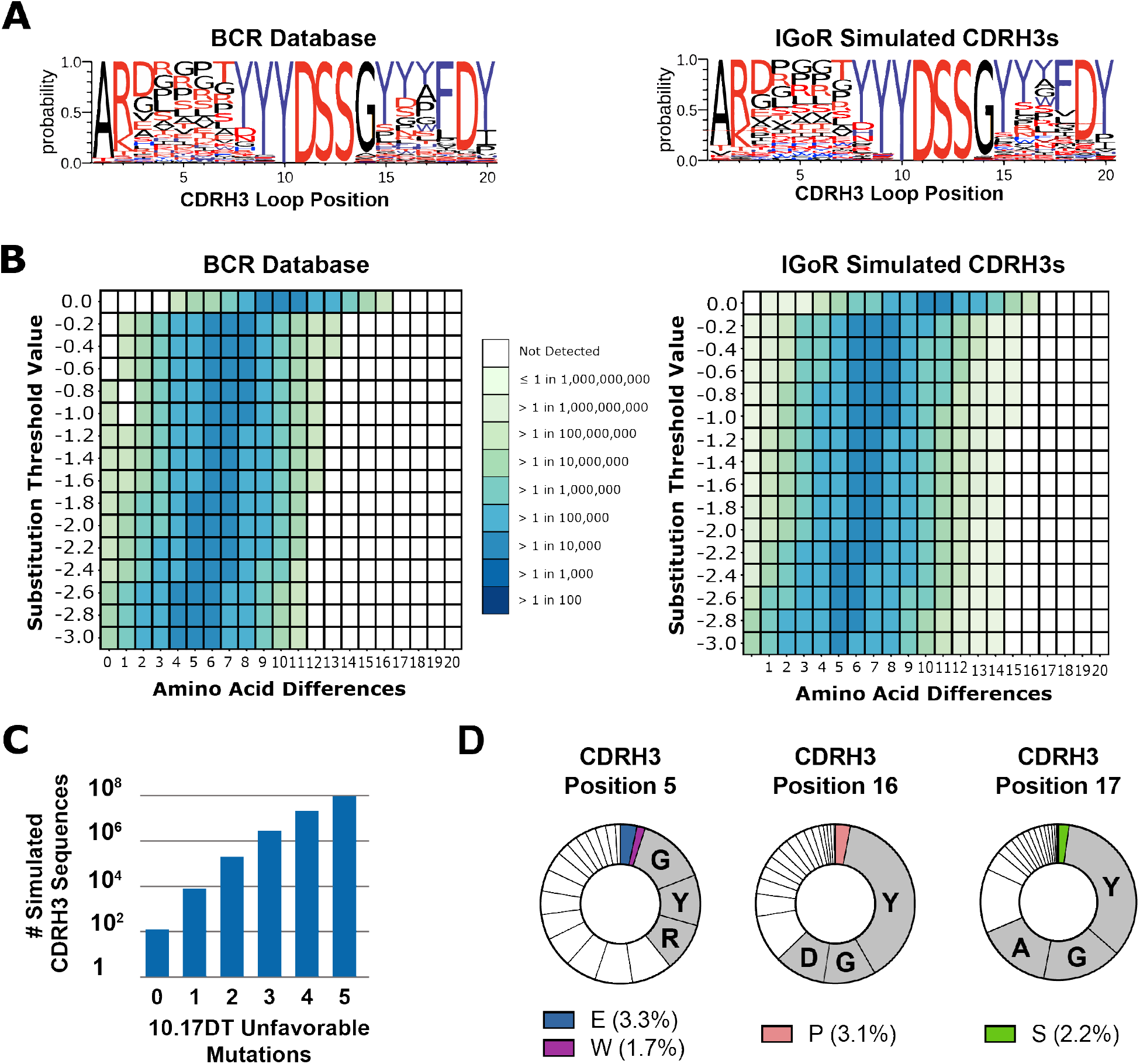
The frequency of CDRH3 loops from the immune repertoire predicted to bind 10.17DT SOSIP. **A.** Sequence logos of DH270 UCA-like CDRH3s loops from either the human BCR database (*left*) or IGoR simulated (*right*). DH270 UCA-like CDRH3s loops contain 20 amino acids and utilize the D3-22 gene with the core “YDSS” motif starting at position 10. Non-polar residues are shown in *black*, polar residues in *red* and aromatic residues in *blue*. **B.** Heatmaps displaying the frequency of 10.17DT SOSIP compatible CDRH3 loops in the experimental BCR sequence database (*left*) and simulated by IGoR (*right*). Each heatmap shows the calculated CDRH3 loop frequency in the respective data set at a given number of amino acid differences (x-axis) and allowing for varying degrees of enrichment threshold values (y-axis) relative to the sequence of the DH270 UCA CDRH3. Enrichment threshold values were determined by deep scanning mutagenesis in **Figure 2B**. **C.** The number of DH270 UCA-like CDRH3 loop sequences simulated with IGoR and binned by their number of calculated unfavorable mutations for 10.17DT SOSIP binding. **D.** The frequency of amino acid at the indicated position in the CDRH3s loops of DH270 UCA-like mAbs present in the experimental BCR database. Residues that 10.17DT SOSIP can recognize are shown in colored wedges; grey wedges indicate the top three most common residues present in DH270 UCA-like CDRH3 loop sequences.

From the ~2.5 trillion IGoR-generated CDRH3 sequences, 1.15 billion had the same length and “YDSS” D gene-derived sequence motif as DH270 UCA. The frequency of simulated sequences that matched these criteria (~1:2,100), was similar to the frequency of sequences with the same properties present in the experimental BCR database (~1:1,800). One hundred and twenty-five simulated CDRH3 loop sequence contained only amino acids predicted to be bound by 10.17DT with high affinity (enrichment score>−0.2), yielding a predicted frequency of B cells that can be engaged by this immunogen *in vivo* of ~1 in 20 billion B cells (**Figure 5B)**.

The clustering heatmaps of IGoR sequences (**Figure 5B**) highlighted that CDRH3 loops compatible with 10.17DT SOSIP binding are rare in the human BCR repertoire. The frequency of compatible CDRH3 loops decays quickly as the number of amino acid differences and enrichment score thresholds are decreased. To generate a strong immune response, it is desirable that a candidate immunogen activates B cells from the natural repertoire at frequencies higher than 1 in 10 million (2). For the BCR database heatmaps, bins with frequencies between 1 in 1 million and 1 in 10 million of the total sequences contained at least 1 amino acid difference at enrichment scores below the −0.2 threshold predictive of 10.17DT binding. Thus, this analysis can inform the design of priming immunogens by revealing how close or how far the candidate immunogen is from engaging target CDRH3 loops that can be found at desirable precursor frequencies. Given that there was at least one amino acid difference in all the CDRH3 loops found at suitable precursor frequencies, we next sought to identify which specific CDRH3 positions were most responsible for limiting the number of BCRs compatible with 10.17DT binding. We computed the frequency of all the amino acids at each position of the DH270-like CDRH3 loops from the database (20 amino acids long with the “YDSS” sequence at position 10 corresponding to D3-22) (**Supplementary Figure 8**). Based on the deep scanning mutagenesis data, 10.17DT recognized less than 5% of the amino acids present in natural CDRH3 loops at positions 5, 16, and 17 (**Figure 2C**). This information can guide immunogen design towards molecules with increased recognition of human BCRs. For example, an improved 10.17DT SOSIP version that could bind CDRH3 loops containing the top three most frequent amino acids found in the BCR database at positions 5, 16, and 17 would increase the frequency of recognized CDRH3s in the repertoire by a factor of ~2300 over 10.17DT (**Figure 5D**). Such an improved immunogen would subsequently increase the estimated frequency of engaged CDRH3 loops from 1 in 20 billion to 1 in 8.5 million, bringing the precursor frequency of the targeted B cells into a range that is more likely to result in robust immune activation (47).

The human BCR database was also analyzed for CDRH3 loop sequences related to CH235 UCA that can be expected to be recognized by the target immunogen CH505.M5.G458Y. In contrast to DH270 UCA, the composition of the CDRH3 sites encoded by the D gene and recognized by CH505.M5.G458Y was not restricted in CH235 UCA (**Figure 3C**). Therefore, we did not limit the BCR database search using any D gene position or composition criteria as we did for DH270 UCA. The database search yielded 9,623,490 CDRH3s that matched the CH235 UCA CDRH3 length of 15 amino acids (11.3% of database sequences). CH505.M5.G458Y binding to CH235 UCA was not affected by the majority of the single amino acid substitutions in the CDRH3 loop (**Figure 3B**). Accordingly, we found that CDRH3 loops predicted to be bound by CH505.M5.G458Y (0 mismatches according to the substitution profile) occurred with a high frequency of 1 in 100 sequences from the human BCR database (**Supplementary Figure 6**).

Overall, these data revealed that the naive B cell repertoire contains a high number of BCRs with CDRH3 loop sequence that permit engagement by CH505.M5.G458Y, while B cell activation by the 10.17DT immunogen will be significantly limited by the lack of BCRs containing CDRH3 loops expected to be bound by this immunogen. By analyzing a large collection of BCRs either sequenced from the naïve B cell repertoire or generated computationally, these methods can rapidly estimate the frequency of characteristic B cells that can be engaged by an immunogen upon vaccination.

## Discussion

To successfully elicit target antibodies by vaccination, a candidate immunogen needs to potently activate related B cell lineages that can subsequently mature through somatic mutations. It is particularly important to explicitly consider precursor engagement during the immunogen design process if target B cells are expected to be rare in the naive repertoire. Indeed, the precursors of some HIV bnAb lineages were estimated to be engaged by candidate immunogens with a frequency as low as ~1 in 50 million B cells (13). Many bnAbs of interest for HIV-1, influenza or coronavirus vaccine development contain rare features such as long CDRH3 loops that are responsible for 50% or more of the binding interaction with the virus (**Supplementary Table 1**) (4–14). Given that CDRH3s loops of over 20 amino acids are present in the natural repertoire in ~13% of the B cells (36, 45), the amino acid composition of the CDRH3 sequence, and not the CDRH3 length, is the primary driver of low precursor frequency. In order to robustly activate a large number of B cells upon vaccination, a candidate immunogen must therefore tolerate multiple amino acids in the CDRH3 loops of target BCRs, particularly in regions generated stochastically by N-nucleotide addition. This is because the overall CDRH3 frequency in the immune repertoire can be estimated as the product of frequencies of tolerated amino acids at each individual position in the CDRH3 loop. Therefore, each additional CDRH3 position in which immunogen recognition is restricted to only a few amino acids leads to a rapid reduction in the overall frequency of natural CDRH3 loops that can be engaged by vaccination.

Typically, the frequency of B cells that can be activated by a given immunogen is estimated by FACS analysis of B cells isolated from human donors (33, 48). Cells are selected based on their ability to selectively bind a target immunogen and their BCRs are subsequently identified and characterized. This approach is laborious, expensive, may be biased by the particular cell labeling and sorting strategy, and requires access to large number of isolated B cells, especially when the target BCRs are expected to be present at low frequency. In comparison, the approach described here relies on bioinformatic analysis of human BCR sequences from either publicly available databases or simulated from models of V(D)J recombination, using sequence identification criteria determined experimentally. For a given antibody-antigen pair, we first employed deep mutational scanning to determine the single amino acid variants in the CDRH3 loop that maintained antigen binding. Based on the resulting data, we then found human BCRs containing CDRH3 loops related to the target antibody and that are expected to be recognized by a particular immunogen. This method allows the identification of CDRH3 loops that are significantly different in sequence than that of the target antibody yet are still expected to be bound by the given immunogen.

Analysis of the CH505.M5.G458Y immunogen found that BCRs containing CDRH3s predicted to bind it are abundant (1:100) in the natural repertoire. Therefore, we expect that vaccination with CH505.M5.G458Y will engage and activate CH235.12 precursors with diverse CDRH3 loops. In contrast, 10.17DT immunogen binding to target CDRH3 loops was highly dependent on amino acid sequence identity. Using traditional immunogenetics definitions of CDRH3 loop length and D gene motif position, DH270 precursors are predicted to occur at high frequency in the natural repertoire (~1:1700). However, library screening of the DH270 UCA antibody clearly showed that 10.17DT recognition of precursor CDRH3 loops is dramatically limited by amino acid composition in N-nucleotide encoded regions. Of the nine amino acid sites inferred to be derived from N-nucleotide additions in DH270, 10.17DT recognizes only 4.2 amino acids on average. This example emphasizes that B cell frequency definitions relying simply on CDRH3 immunogenetic features like length, D gene usage and D gene position without regard to the specific amino acid composition of the CDRH3 could lead to gross overestimations of the frequencies of B cells that can be engaged by candidate immunogens.

To sample the immune repertoire at sequence depths beyond those available in databases of isolated human BCRs, we demonstrated that IGoR simulations can accurately model natural CDRH3 loop sequence diversity. These synthetic sequences can then be analyzed to determine the frequency of rare BCRs that may be engaged by a target immunogen. We were able to use these synthetic sequences to generate a realistic precursor frequency estimate for DH270-like BCRs that could be engaged by 10.17DT that could not have been otherwise obtained through sequence repository searches. Additionally, this bioinformatic analysis can be used to inform an immunogen design strategy to optimize precursor engagement. By searching for compatible CDRH3 sequences in large repertoire sequence databases or by simulating V(D)J recombination, this approach can precisely identify features limiting precursor frequency for a particular immunogen which gives actionable design information for how to improve diversity of B cell precursor engagement by targeting more frequent amino acids in the BCR repertoire at specific positions of the CDRH3. As an example of how to apply this information to priming immunogen design, we showed that modifying the 10.17DT immunogen to accommodate the three most frequent amino acids at three additional positions would be predicted to increase the precursor frequency by over 3 orders of magnitude.

Despite the advantages over B cell sorting and isolation, the CDRH3 centric approach used in this study has some limitations in estimating the frequency of B cells targeted by a given immunogen. It is possible that some CDRH3 loops identified as favorable by our method could be part of BCRs that contain other molecular features that prevent immunogen binding, such as incompatible VH or VL gene segments (13, 22, 49). In addition, the deep scanning mutagenesis approach evaluates the effect on immunogen binding of only one amino acid change in the CDRH3 loop of target antibodies. However, as shown in this study, many of the BCRs identified contain multiple amino acids changes relative to the target CDRH3. These loops may contain novel structural and molecular features where the additive effect of individual mutations may no longer predict the overall binding propensity to the target immunogen. Nevertheless, in this study we were able to identify and validate experimentally CDRH3 loops that showed binding to 10.17DT yet had up to 45% different amino acids than those of the DH270 UCA CDRH3. Our analysis was additionally constrained to sequences with matching CDRH3 lengths; however, it is possible that immunogens may recognize BCRs that contain CDRH3 loops of different lengths that could affinity mature into bnAbs either by acquiring indels or through presently unknown evolutionary pathways.

While in this study we focused our analysis on naïve B cell receptor CDRH3 loop sequences, our method is generalizable. Other BCR features important for immunogen recognition, such as variable heavy or light chain gene templates or specific acquired somatic mutations, could be used as search criteria. We believe this combined experimental and bioinformatic method will prove to be a valuable tool for immunogen characterization and optimization.

## Author Contribution

OS, JSMB, BFH, KW and MLA developed the study and designed experiments; OS, BR, AW and MLA performed and analyzed the deep scanning mutagenesis experiments; OS, BR, MB and RP performed binding experiments of SOSIPs to recombinant antibodies; JSMB and KW performed the computational analysis of the BCR database and IGoR simulations; HC and KOS provided SOSIP molecules; BFH, KW and MLA acquired the funding to support this work; OS, JSMB, KW and MLA wrote the first draft of the manuscript and the other authors provided edits.

## Acknowledgements

We would like to thank Sravani Venkatayogi for bioinformatics support. We would like to thank Duke University and OIT Research Computing for providing computational resources and data storage through the Duke Compute Cluster. We thank the staff of the Viral Genetics and Analysis Core Facility in the Duke Human Vaccine Institute for help with library prep and Illumina sequencing. Fluorescence Activated Cell Sorting of yeast display libraries was performed at The Flow Cytometry Shared Resource (FCSR) in the Duke Cancer Institute and the Duke Human Vaccine Institute Research Flow Cytometry Facility. Surface Plasmon Resonance measurements were carried at the Duke Human Vaccine Institute Biomolecular Interaction Analysis Facility.

## Data availability

The code used for bioinformatics analysis is deposited and publicly available from GitHub (https://github.com/WieheLab/CDRH3_ScanPan); other data that support the findings of this study and that are not described in the main text or the supplementary material are available from the corresponding authors (KW and MLA).

## Methods

### Plasmids and DNA synthesis

CDRH3 single site mutagenesis libraries were synthesized on a BioXp system (CodexDNA). For CH235 UCA CDRH3 libraries, heavy chain residues 95-107 encoded by the D gene, the N-nucleotide addition, as well as the first three amino acids of the JH gene (amino acid residues 95-107) were allowed to sample all the 20 amino acids. For DH270 UCA CDRH3 libraries, the parent sequence DH270UCA3 was utilized as the base template and heavy chain residues 97-116 encoded by the last two amino acids of the VH gene, the D gene, the N-nucleotide additions, and the first three residues of the JH gene were allowed to sample all the 20 amino acids. Genes encoding the antibody heavy and light chains were commercially synthesized and cloned into pcDNA3.1 vector (GenScript). DNA primers for sequencing and insert amplification were ordered from IDT.

### Development and screening of scFv libraries on the surface of yeast

#### Library design and synthesis

Single site saturation mutagenesis libraries were synthesized as described above. Library scFvs were then amplified with Q5 polymerase and purified by agarose gel extraction and PCR cleanup (Qiagen) as per the manufacturer’s protocol.

#### Library transformation into S. cerevisiae

*S. cerevisiae* EBY100 yeast cells were transformed to express the CDRH3 single site saturation mutagenesis libraries as previously described(42, 50, 51). Cells were transformed by electroporation with a 3:1 ratio of 12μg scFv library DNA and 4μg pCTCON2 plasmid digested with BamHI, SalI, NheI (NEB). The typical sizes of the transformed libraries, determined by serial dilution on selective plates, ranged from 2-6×10^7^. Between 60% and 80% of the sequences recovered from the transformed libraries were confirmed to contain full length, in-frame genes by Sanger sequencing (Genewiz). Yeast libraries were grown in SDCAA media (Teknova) supplemented with pen-strep at 30C and shaking (225 rpm).

#### Library screening by FACS

scFv expression on the surface of yeast was induced by culturing the libraries in SGCAA (Teknova) media at a density of 1×10^7^ cells/mL for 24-36 hours. Cells were washed twice in ice cold PBSA (0.01M sodium phosphate, pH 7.4, 0.137M sodium chloride, 1g/L bovine serum albumin) and incubated for 1 hour at 4C with 300nM biotinylated immunogens expressed as HIV-1 Env SOSIPs (10.17DT for the DH270 library and CH505_M5_G458Y for the CH235 library). Cells were then washed twice with PBSA, resuspended in secondary labeling reagent α c-myc:FITC (ICL) and streptavidin:R-PE (Sigma-Aldrich) and incubated at 4C for 30 minutes. Cells were washed twice with PBSA after incubation with the fluorescently labeled probes and sorted on a FACS-DiVa (BD). Double positive cells for PE and FITC were collected and expanded for one week in SDCAA media supplemented with pen-strep before DNA isolation from selected clones. FACS data was analyzed with Flowjo_v10.6 software (Becton, Dickinson & Company). All clones selected by FACS were expanded, and their DNA was extracted (Zymo Research) for analysis by Next Generation Sequencing (Illumina) and Sanger sequencing (Genewiz).

### Sequence analysis of isolated library clones

ScFv encoding plasmids were recovered from yeast cultures by miniprep with the Zymoprep yeast plasmid miniprep II kit (Zymo Research) as previously described(42). Isolated DNA was transformed into NEB5α *E. coli* (NEB), and the DNA of individual bacterial colonies was isolated (Wizard Plus SV Minipreps, Promega) and analyzed by Sanger sequencing. To prepare for Next Generation Sequencing, the scFv insert from isolated plasmids was amplified by PCR using Q5 polymerase (NEB). DNA samples were prepped and run using the Illumina MiSeq v3 reagent kit following manufacturer’s protocols. Illumina sequencing returned an average of 21.6 million reads per sample, of which an average of 20.7 million mapped to the scFv amplicon. Sequencing data was processed using Geneious Prime and in-house scripts to compute the amino acid frequency and distribution.

### Natural human BCR repertoire analysis

We downloaded locally the BCR sequence database developed by Briney et al., which was constructed from deep sequencing 10 individuals’ BCR repertoires (36). We deduplicated all VDJ sequences per subject and discarded any VDJ sequences that contained premature stop codons. The resulting dataset was comprised of 85,149,053 sequences. Frequency matrices of the naïve (N) and sorted experimental libraries (S) were created using the amino acid counts for each position as determined by NGS sequencing. A log fold change matrix was subsequently calculated using F_ij_=log_2_(S_ij_/N_ij_) for each substitution *i* at each position *j*. The curated BCR sequence database was queried first for sequences with the same CDRH3 length, same D gene, D gene reading frame and position in the case of DH270 UCA, or by same CDRH3 length in the case of CH235 UCA. Sequences that met these criteria were then measured against the CDRH3 amino acid substitution profile of their target antibody by considering any amino acid with a log fold change of −0.2 or higher as a match (Supplementary Figure 3). Additionally, heatmaps were generated by varying the binding thresholds used to define a match/mismatch and computing the frequency of sequences within the input dataset that occurred over a range of distances from 0 to N mismatches where N is the length of the CDRH3. For production of chimeric DH270 UCA antibodies with the most DH270-like CDRH3s from the BCR database, we first selected CDRH3 sequences with the same length and then scored each sequence by summing the inverse of the fold change value over each position in the CDRH3 and selecting the top 100 scoring CDRH3s for experimental characterization.

#### CDRH3 sequence simulation

We employed the IGoR program to simulate VDJ recombination and generate synthetic CDRH3 sequences(38). VDJ sequences were generated without using the IGoR error model and thus represent unmutated sequences. As sequences were generated, we stored only CDRH3s that matched the criteria for being DH270 UCA-like: same length and with the “YDSS” motif at position 10 indicative of the germline gene D3-22. This resulted in 5×10^12^ total generated sequences with 3.3×10^11^ CDRH3 sequences of the same length as DH270 UCA and 1.15×10^9^ sequences with the same length and D-gene representative YDSS motif located at position 10 of the CDRH3.

### Antibody expression and purification

Antibodies were expressed and purified as previously described(11, 42). Briefly, 100mL cultures of Expi293F cells at a density of 2.5×10^6^ cells/mL were transiently transfected with 50μg each heavy and light chain encoding plasmids and Expifectamine (Invitrogen) per manufacturer’s protocol. Five days after transfection, cell culture media was cleared of cells by centrifugation, and the supernatant was filtered through a 0.8μm filter (Nalgene). Clarified supernatant was incubated with Protein A beads (ThermoFisher) over night at 4C, washed with 20mM Tris supplemented with 350mM NaCl (pH=7), followed by elution with a 2.5% Glacial Acetic Acid Elution Buffer and subsequent buffer exchange into 25mM Citric Acid supplemented with125mM NaCl (pH=6). IgGs expression was confirmed by reducing SDS-PAGE analysis and quantified by measuring absorbance at 280nM (Nanodrop 2000).

### Recombinant HIV-1 Env SOSIP production

SOSIP envelopes were produced recombinantly as previously described (11). Briefly, Freestyle 293 cells were transfected with 293Fectin complexed with envelope-expressing DNA and furin-expressing plasmid DNA. After 6 days, SOSIPS were purified via PGT145 affinity chromatography with subsequent size exclusion chromatography. Trimeric HIV-1 Env fractions were pooled, flash-frozen, and stored at −80C in 10mM Tris pH 8, 500mM NaCl buffer.

### Binding analysis by Surface Plasmon Resonance

10.17DT binding of DH270 UCA antibody variants containing single site mutations (Figure 2E) as binding of DH270 UCA chimeric antibodies containing CDRH3 loops identified from the natural BCR database (Figure 4) was analyzed by Surface Plasmon Resonance on a BIAcore 3000 instrument (GE Healthcare). Biotinylated SOSIP Env 10.17DT was immobilized on a CM5 chip coated with streptavidin. Antibodies were then injected at different concentrations to determine the maximum (Rmax) binding response to 10.17DT. Background binding levels to 10.17DT obtained by buffer alone and by a control non-HIV antibody (palivizumab) were subtracted from the measured response. For measurements of the single mutant DH270 UCA antibodies (Figure 2E), 3000-4000 RUs of 10.17DT The 10.17DT SOSIP was amine coupled on the sensor chip and mutated DH270 UCA antibodies were injected at 100nM to assess their binding avidity relative to that of the wild-type antibody. For measurements of the DH270 UCA chimeric antibodies (Figure 4), 2000-3000 RUs of 10.17DT were captured and antibodies were injected at concentrations of 20μM and 2μM. Data to determine the Rmax value was analyzed using the BIAevaluation 4.1 software (GE Healthcare). Rmax values were reported as relative signal to that obtained by the interaction of WT DH270 UCA antibody to 10.17DT.

### Binding analysis by ELISA

Binding to 10.17DT by DH270 UCA single mutant antibodies at position L104 (Supplementary Figure 4) and by a subset of DH270 UCA antibody variants containing single site mutations (Figure 2E) was measured by ELISA. The target antibody was captured on 96 well plates overnight, and then incubated with serial dilutions (1:3) of 10.17DT SOSIP, starting at a concentration of 100μg/mL. Binding was detected by incubation with biotinylated antibody PGT151, that is specific Env trimers, followed by addition of streptavidin-HRP. Plates were read at 450nm on a SpectraMax 384 PLUS reader (Molecular Devices). The logarithm of the area under the curve (LogAUC) was calculated using Prism 9.

## Supplementary Figures

**Supplementary Table 1.** Multiple antibodies against diverse viruses contain long CDRH3 loops that are essential for viral recognition. Antibody/antigen structural complexes describing a wide variety of antibody bound to viral proteins were analyzed using PDBePISA to determine the buried solvent accessible surface area (BSASA) at the binding interface in units of Å^2^. BSASA binding contributions for residues in the light chain (LC) heavy chain (HC), and CDRH3 loop are reported for each antibody.

| Antibody | Virus | Viral Protein Target | PDB ID | CDRH3 Length | HC BSASA | LC BSASA | BSASA Total | BSASA CDRH3 | % BSASA CDRH3 |
| --- | --- | --- | --- | --- | --- | --- | --- | --- | --- |
| **EDE2 A11** | Dengue/Zika | E | 4UTB | 26 | 512 | 59 | 571 | 438 | 77% |
| **HEPC3** | HCV | E2 | 6MEI | 19 | 953 | 68 | 1022 | 580 | 57% |
| **CH235** | HIV | Env CD4 bs | 6UDA | 13 | 613 | 275 | 889 | 275 | 31% |
| **CH03** | HIV | Env V1V2 | 5ESV | 26 | 385 | 0 | 385 | 263 | 68% |
| **VRC26.25** | HIV | Env V1V2 | 6VTT | 38 | 1115 | 0 | 1115 | 1115 | 100% |
| **BG18** | HIV | Env V3 | 6DFH | 23 | 395 | 326 | 721 | 389 | 54% |
| **DH270** | HIV | Env V3 | 6UM5 | 20 | 410 | 59 | 469 | 242 | 52% |
| **CH65** | Influenza | HA (RBS) | 5UGY | 19 | 554 | 296 | 850 | 396 | 47% |
| **FI6V3** | Influenza | HA (stem) | 3ZTJ | 22 | 542 | 320 | 863 | 532 | 62% |
| **1G01** | Influenza | NA | 6Q23 | 21 | 780 | 251 | 1031 | 841 | 82% |
| **DH1047** | SRA-CoV-2 | RBD | 7Ld1 | 24 | 546 | 238 | 784 | 380 | 48% |

**Supplementary Figure 1.**
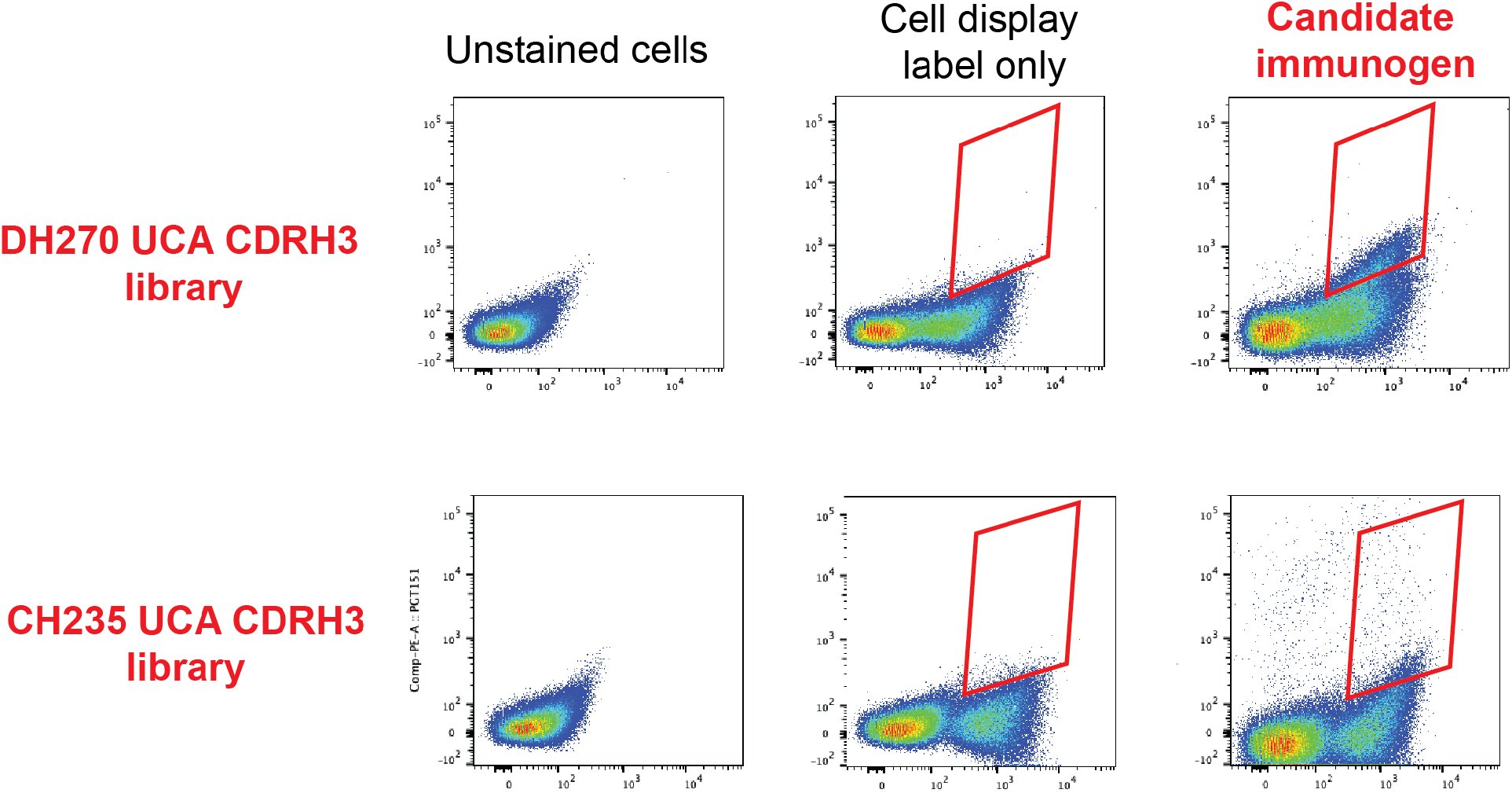
FACS selection of DH270 UCA and CH235 UCA CDRH3 single-site saturation libraries for binding to the target immunogens. 10.17DT and CH505.M5.G458Y scFv libraries were displayed on the surface of yeast and labeled for cell surface display (x-axis) and binding to the candidate immunogen (y-axis). DH270 UCA and CH235 UCA libraries were sorted with 300nM concentration of biotinylated SOSIP target immunogens 10.17DT and CH505.M5.G458Y respectively, and all binding clones were recovered for downstream DNA isolation and NGS analysis.

**Supplementary Figure 2.**
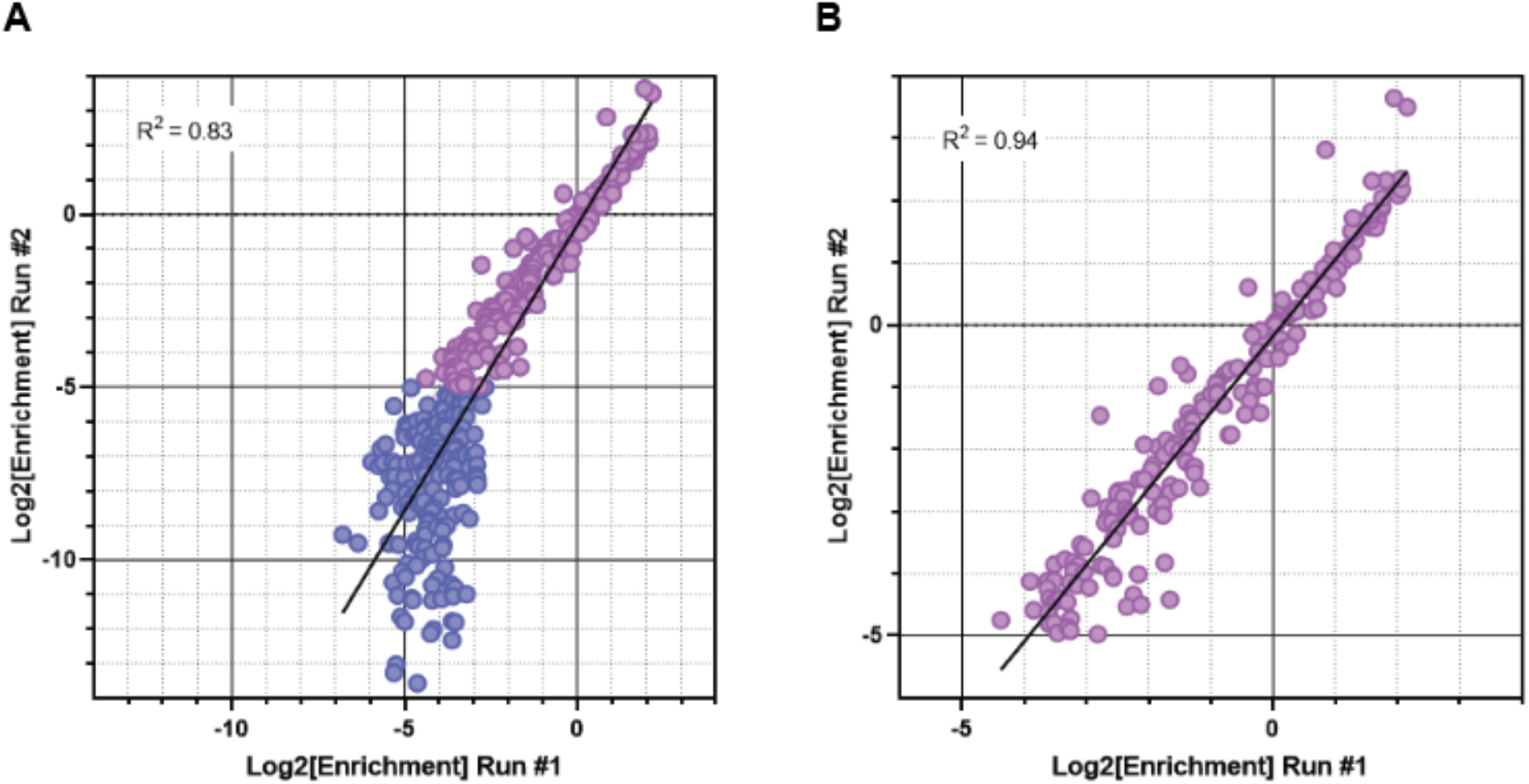
Independent sort experiments generate strongly correlated enrichment scores. **A.** Enrichment scores of two sort replicates completed on separate day are shown. The correlation was strongest for enrichment scores higher than −5 (pink) and decreased somewhat for values below this threshold (purple). An enrichment score of 5 corresponds to a 32-fold depletion of a particular mutation from the initial library upon sorting with 10.17DT.

**Supplementary Figure 3.**
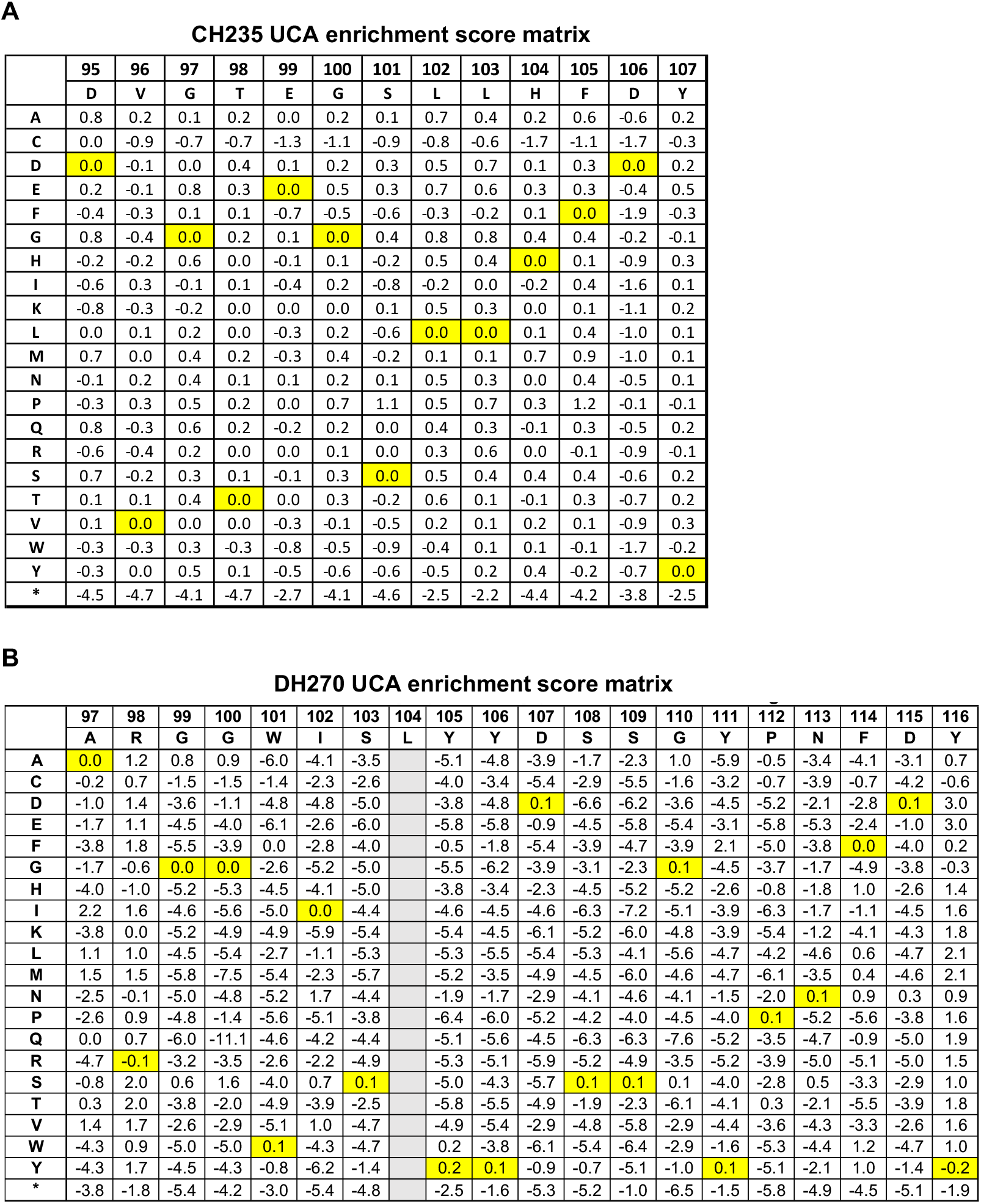
scFv library enrichment score matrices. Enrichment scores, calculated as log_2_ (fold change) of scFv residues before and after sorting with candidate immunogens. **A.** Enrichment scores for each position of CH235 UCA CDRH3 after selection with immunogen CH505.M5.G458Y. **B.** Enrichment scores for each position of DH270 UCA CDRH3 after selection with immunogen 10.17DT. Wild type residues shown in yellow. * Stop codon.

**Supplementary Figure 4.**
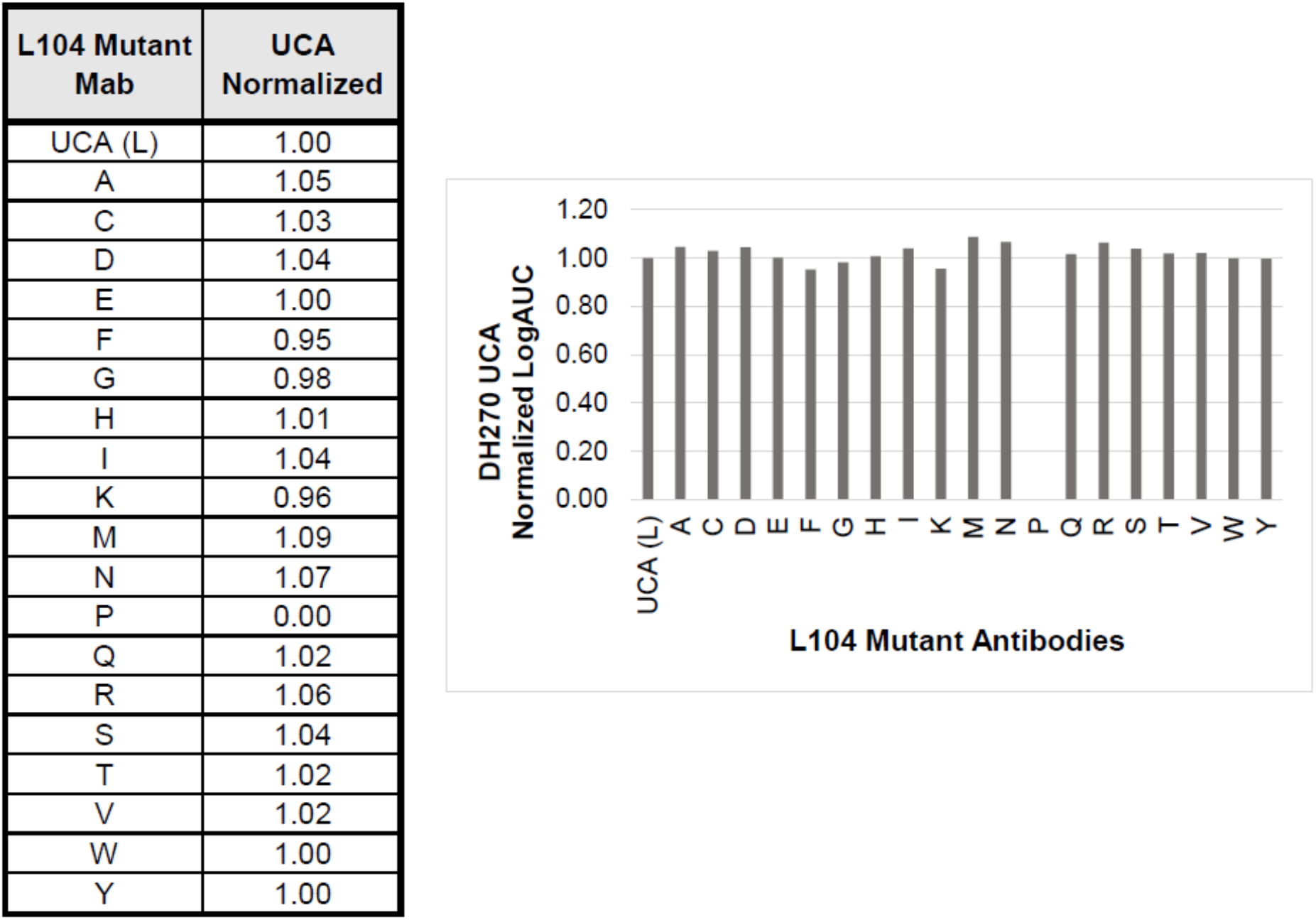
Binding characterization of DH270 UCA mutant antibody variants at position 104. Single site mutated antibodies containing every possible substitution residue at position 104 of DH270 UCA were recombinantly produced and tested for binding to 10.17DT target immunogen. Results are reported as logAUC, normalized to DH270 UCA binding signal.

**Supplementary Figure 5.**
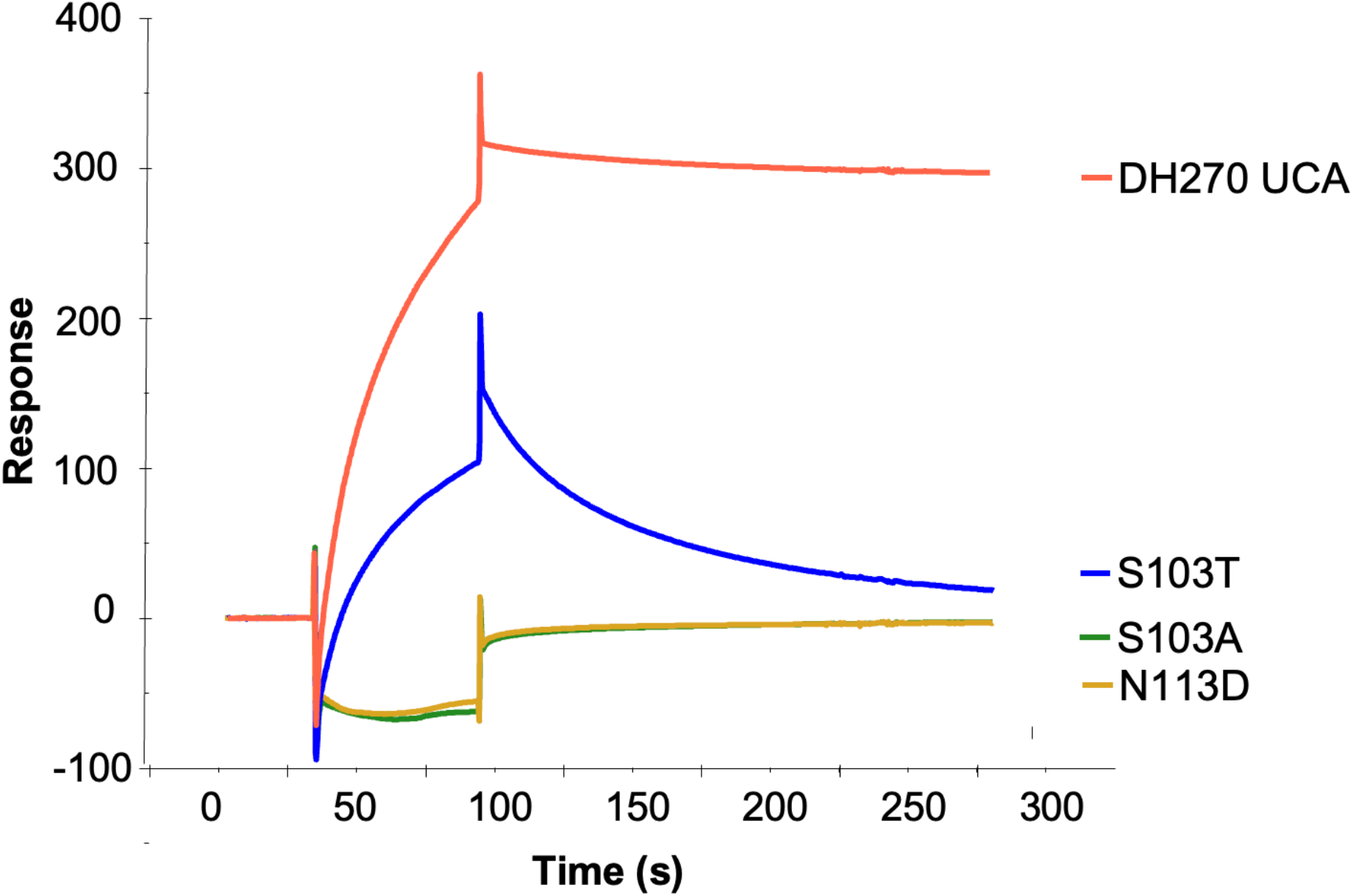
Surface Plasmon Resonance of DH270UCA antibodies with point mutants predicted to decrease 10.17DT binding.

**Supplementary Figure 6.**
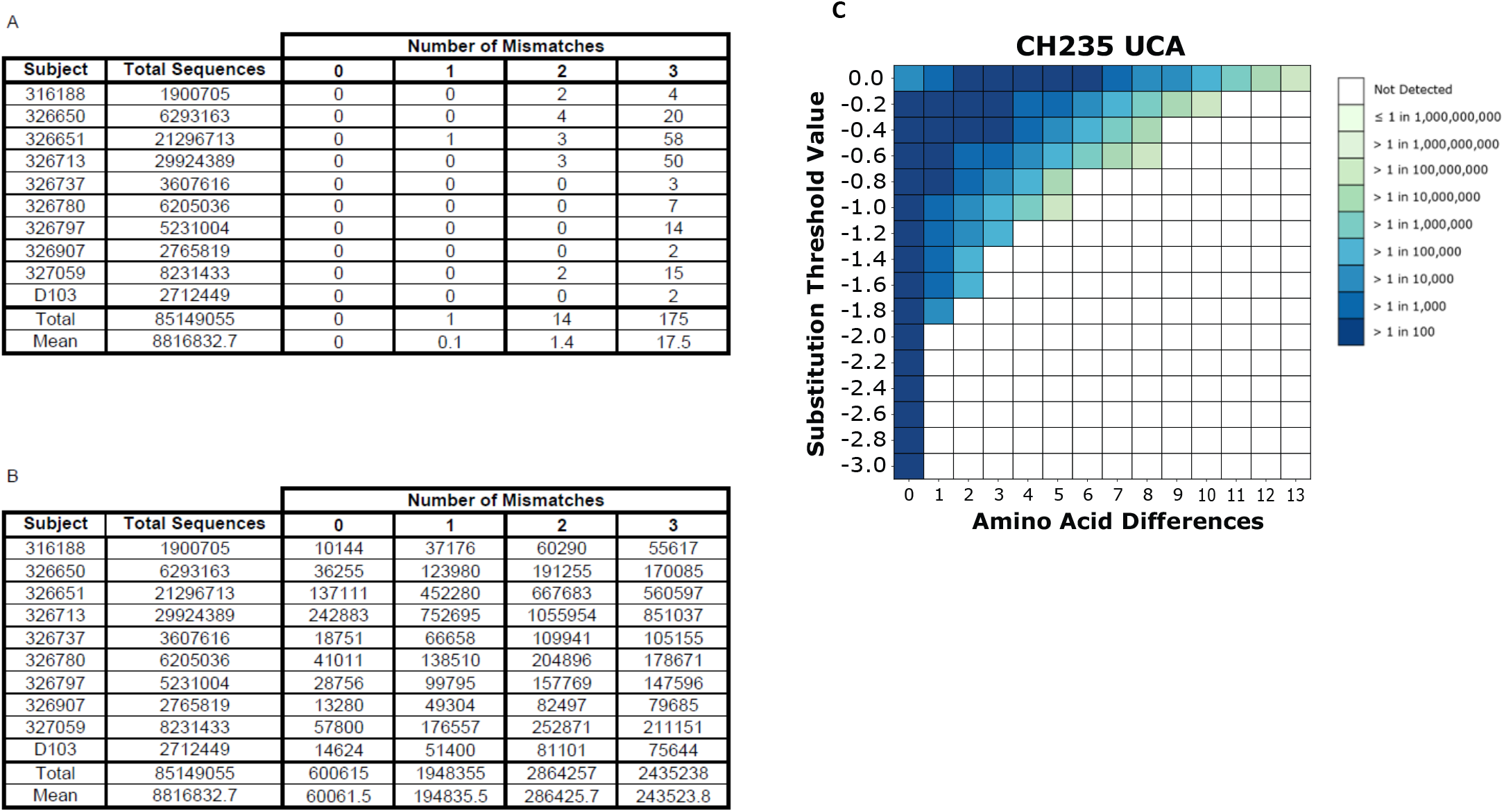
Number of natural CDRH3 predicted to be recognized by the candidate immunogens. **A.** The total number of sequences isolated together with the number of CDRH3 sequences that contain 0, 1, 2, or 3 mismatches relative to the DH270 UCA CDRH3 substitution profile recognized by 10.17DT. **B.** The total number of sequences isolated together with the number of CDRH3 sequences that contain 0, 1, 2, or 3 mismatches relative to the CH235 UCA CDRH3 substitution profile recognized by CH505.M5.G458Y. **C.** CH505.M5.G458Y compatible (CH235 UCA-like) CDRH3s in experimental BCR sequence database. Heatmap shows the calculated CDRH3 loop frequency for a given number of amino acid differences (*x-axis*) and allowing for varying degrees of substitution threshold values (*y-axis*) relative to the sequence of CH235 UCA. Substitution threshold values were determined by deep scanning mutagenesis in **Figure 3B**. Higher substitution threshold values correspond to a stricter definition of what constitutes a matching amino acid in the CDRH3 substitution profile.

**Supplementary Table 2.**
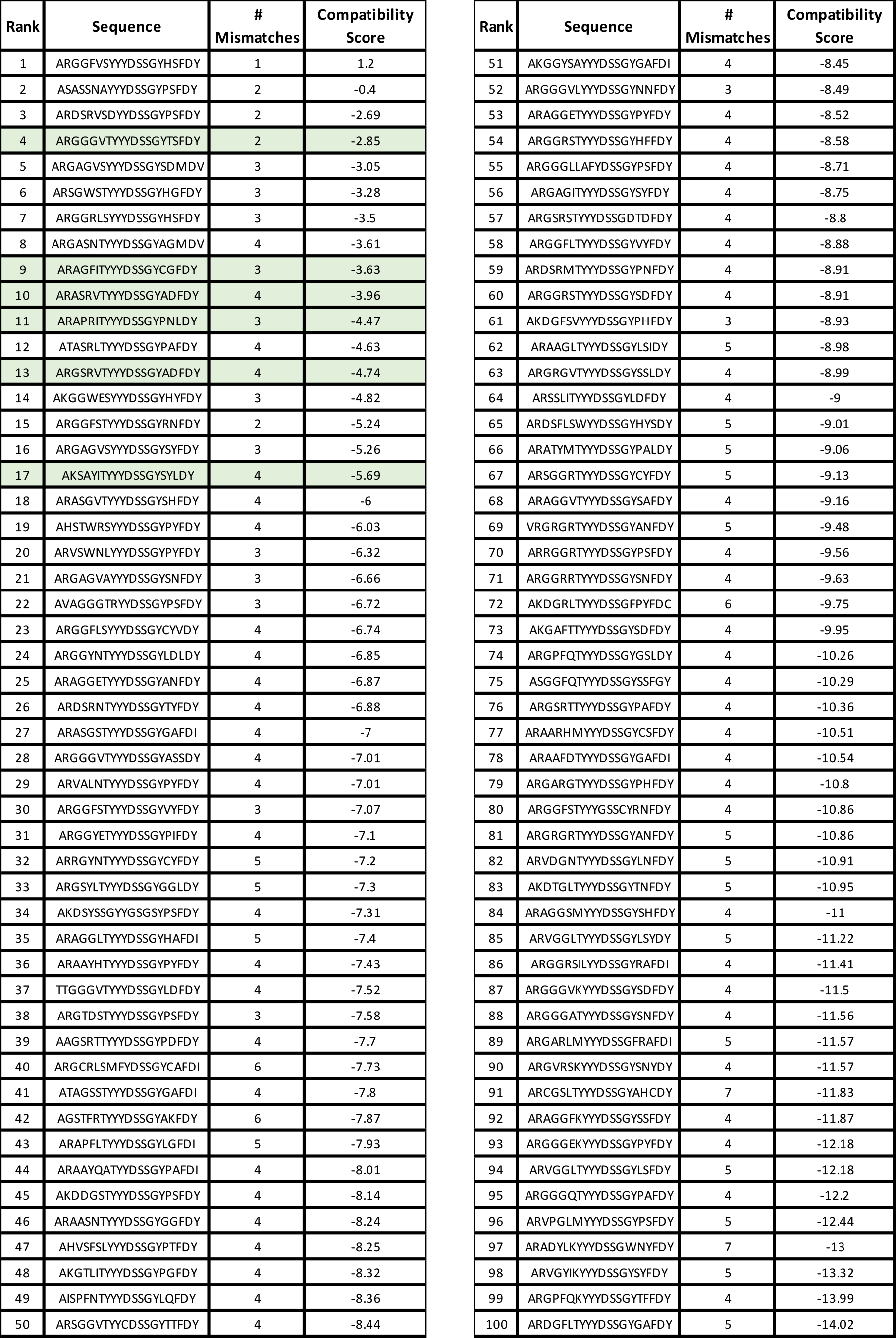
The sequence, number of mismatches, and compatibility scores of 100 natural CDRH3 sequences using the substitution profile of DH270 UCA towards 10.17DT binding. Chimeric DH270 UCA antibodies containing these CDRH3 loops were expressed as an scFv library on the surface of yeast and assayed for binding to 10.17DT immunogen. The number of mismatches is the number of amino acids with an enrichment score of less then −0.2. The compatibility score is the sum of enrichment scores for the entire sequence. Antibodies recombinantly expressed and characterized in **Figure 4** and **Supplementary Figure 7** highlighted in green.

**Supplementary Figure 7.**
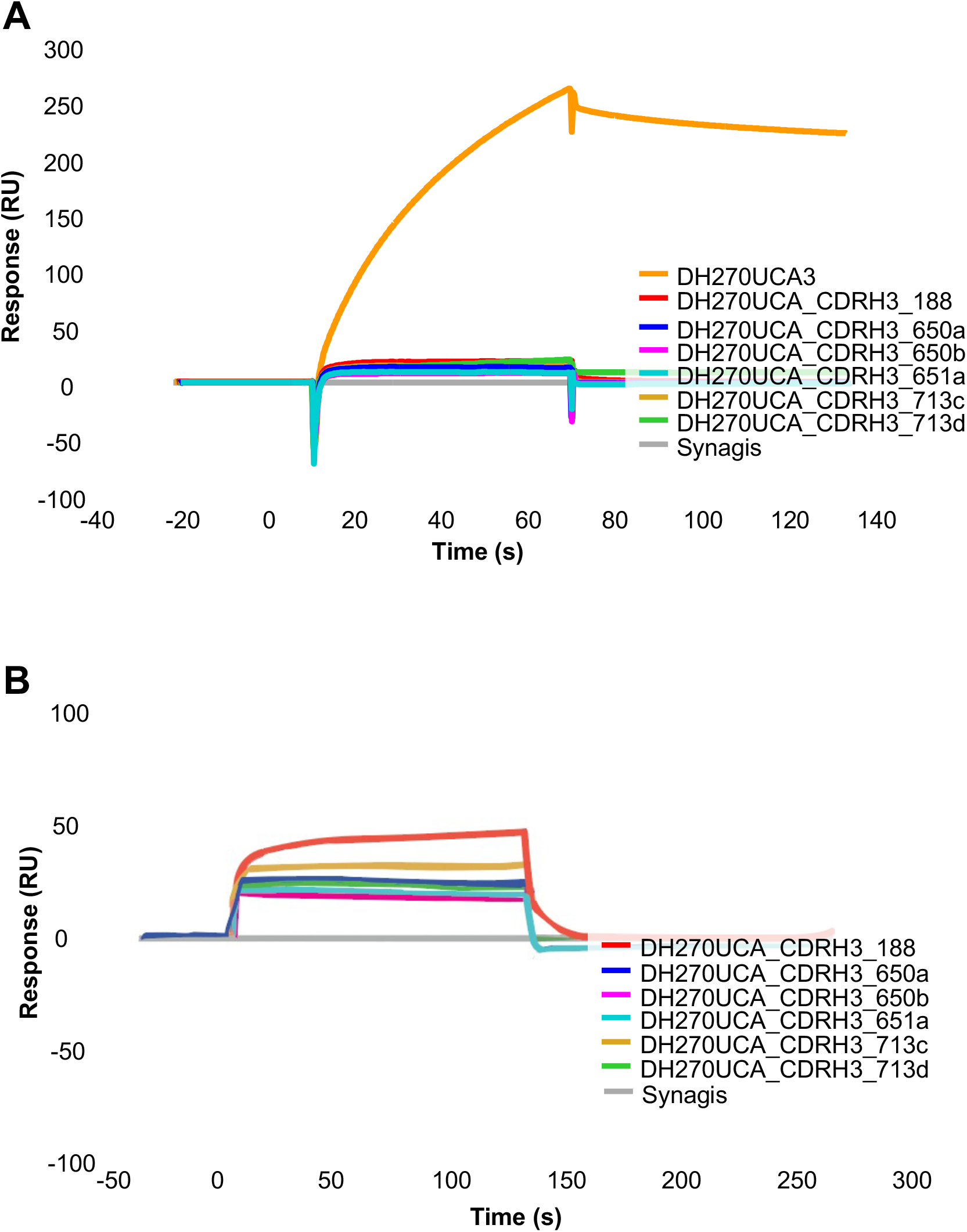
Surface Plasmon Resonance of chimeric DH270 UCA antibodies with natural CDRH3 sequences. **A.** 10.17DT SOSIP immobilized on chip (2784 RU) with monoclonal antibodies injected at 12 μM concentration. **B.** Chip from **A** with chimeric CDRH3 antibodies and negative control (Synagis) injected at concentration of 20 μM.

**Supplementary Figure 8.**
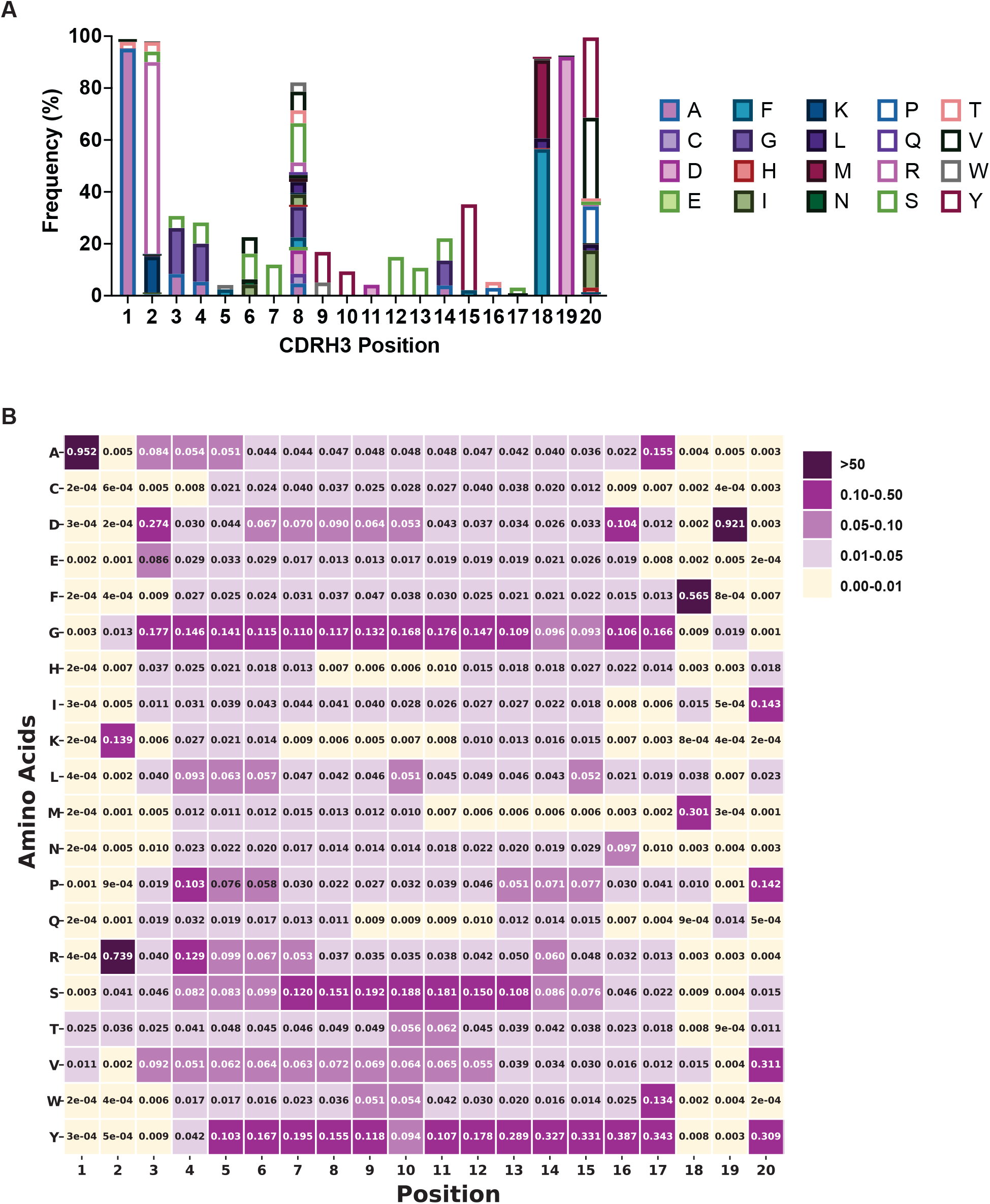
Positional Frequency of 10.17DT Compatible Amino Acids in the BCR Database. **A.** The amino acid frequency at each position amongst 20 amino acid long CDRH3 sequences from the BCR database estimated to be compatible with 10.17DT binding (enrichment score > −0.2) in the DH270 UCA CDRH3 substitution profile. Positions 5, 16, and 17 are highly restricted relative to the amino acids observed in these positions in the BCR database. **B.** The frequency of all amino acids at each position in 20 amino acid long CDRH3s in the BCR database. Immunogens that can bind and tolerate alternative high frequency amino acids at position 5 (e.g. Gly, Tyr, and Ser), at position 16 (e.g. Tyr, Gly, and Asp), and at position 17 (e.g. Tyr, Gly, and Ala) would greatly improve the overall frequency of compatible CDRH3s in the BCR repertoire.

